# Physiological adaptation in flagellar architecture improves *Vibrio alginolyticus* chemotaxis in complex environments

**DOI:** 10.1101/2023.02.06.526967

**Authors:** Marianne Grognot, Jong Woo Nam, Lauren E. Elson, Katja M. Taute

## Abstract

Bacteria navigate natural habitats with a wide range of mechanical properties, from the ocean to the digestive tract and soil, by rotating helical flagella like propellers. Species differ in the number, position, and shape of their flagella, but the adaptive value of these flagellar architectures is unclear. Many species traverse multiple types of environments, such as pathogens inside and outside a host.

We investigate the hypothesis that flagellar architectures mediate environment-specific benefits in the marine pathogen *Vibrio alginolyticus* which exhibits physiological adaptation to the mechanical environment. In addition to its single polar flagellum, the bacterium produces lateral flagella in environments that differ mechanically from water. These are known to facilitate surface motility and attachment. We use high-throughput 3D bacterial tracking to quantify chemotactic performance of both flagellar architectures in three archetypes of mechanical environments relevant to the bacterium’s native habitats: water, polymer solutions, and hydrogels. We reveal that lateral flagella impede chemotaxis in water by lowering the swimming speed but improve chemotaxis in both types of complex environments. Statistical trajectory analysis reveals two distinct underlying behavioral mechanisms: In viscous solutions of the polymer PVP K90, lateral flagella increase the swimming speed. In agar hydrogels, despite lowering the swimming speed, lateral flagella improve overall chemotactic performance by preventing trapping in pores.

Our findings show that lateral flagella are multi-purpose tools with a wide range of applications beyond surfaces. They implicate flagellar architecture as a mediator of environment-specific benefits and point to a rich space of bacterial navigation behaviors in complex environments.

## Introduction

Many bacteria swim by rotating flagella, helical appendages driven by a rotary motor at their base. Species vary widely in their flagellar architectures^1,2^. While many marine bacteria swim with only a single polar flagellum (monopolar flagellation), others, including the well-studied enteric *Escherichia coli*, use variable numbers of flagella distributed all over the cell body (peritrichous flagellation). Motility enables bacteria to perform chemotaxis, that is, biased motion aligned with chemical gradients. While chemotaxis provides an evolutionary benefit by increasing nutrient uptake^3^, the evolutionary forces driving different flagellar architectures are unknown. On the one hand, the monopolar flagellation of marine bacteria has been speculated to reflect the biosynthetic cost of flagella in the face of the nutrient scarcity of the ocean^4^. On the other hand, peritrichously flagellated species were reported to display no chemotactic performance benefit compared to monopolar species^5^.

Intriguingly, several species, including a number of pathogens, display physiological adaptation in flagellar architecture^6^. Many *Vibrios*^7^, *Aeromonas*^8^ and others^9–11^ possess a dual flagellar system: a constitutively expressed polar flagellum driven by a sodium ion gradient across the cell membrane as well as peritrichous lateral flagella driven by a proton gradient that are induced only under specific physical or chemical environmental conditions^12–14^. Flagellation is recognized as a major virulence factor in *Aeromonas* spp.^15^ and *Vibrio* spp.^16^, and lateral flagella have been demonstrated to be a host colonization factor in *Vibrio parahaemolyticus*^17^. A dual flagellar architecture has also recently been described in other pathogenic species such as *Plesiomonas shigelloides*^9^ and *Burkholderia dolosa*^18^, isolated from clinical samples.

The established paradigm is that each flagellar system has a distinct role^11^: the polar flagellum drives swimming motility, while the role of lateral flagella is confined to surfaces, enabling swarming motility^11,19,20^ and attachment to surfaces such as the host epithelium^21^. Yet it has long been recognized that lateral flagella are expressed not only upon surface contact and swarming, but under a broad range of conditions that impede the rotation of the polar flagellum, ranging from growth in hydrogels or viscous solutions^12,22^, to agglutination with flagellum-targeting antibodies^12^ as well as inhibition of the polar flagellar motor by a sodium channel inhibitor^13^. Few studies have ventured beyond the functional dichotomy of bulk liquid versus surface use for the two phenotypes of the dual flagellar architecture. Atsumi *et al*.^23^ reported that lateral flagella increase the average swimming speed of *Vibrio alginolyticus* in viscous media. Bubendorfer *et al*.^24^ showed that lateral flagella can increase the colony expansion rate of *Shewanella putrefaciens* in soft agar plates as well as random spreading in buffer by increasing the cells’ directional persistence during turning events.

The ecology of the marine and estuarine pathogen *V. alginolyticus* implies life in a range of environments beyond the liquid-versus-surface dichotomy. Like its close relative, *V. parahaemolyticus, V. alginolyticus* is a pathogen of marine species^25^ as well as an opportunistic human pathogen, mostly infecting open wounds^25,26^, but also a rising foodborne pathogen^27^. It is also found in coral mucus, both in healthy and diseased states^28,29^. Work on mutants defective in lateral flagella has shown that the polar-only phenotype of *V. alginolyticus* executes a so-called run-reverse-flick motility pattern^5,30^ and exhibits efficient chemotaxis towards rare or transient resources^5,31^.

We hypothesized that the physiological adaptation in the flagellar phenotype to the mechanical environment might mediate environment-specific benefits of the resulting motility or chemotaxis behavior. Which flagellar phenotype is optimal, in which environments? To address such questions, a quantitative metric is required to compare the performance of flagellar phenotypes in different environments. Lateral flagella, like the polar flagellum, are under the control of the chemotaxis signaling system^32^. Chemotaxis has been shown to increase nutrient uptake in marine bacteria^33^, suggesting that chemotactic performance might be selected for in *V. alginolyticus*’s natural ecology. While previous work focused on swimming speed^23^, swarming behavior^22^, and random spreading^24^ of the two flagellar phenotypes, we thus chose to investigate their chemotactic performance in specific, mechanically distinct environments.

Here, we harness a recent multiscale 3D chemotaxis assay^34^ to determine the chemotactic performance of the two flagellar phenotypes of *V. alginolyticus* in a set of archetypical mechanical environments: buffer, polymer solutions, and hydrogels. By 3D tracking tens of thousands of cells navigating chemical gradients, we demonstrate that lateral flagella lower the chemotactic drift speed in buffer but increase it in both archetypes of complex materials. We then reveal strikingly different behavioral mechanisms underlying the chemotactic performance benefits observed in hydrogels and polymer solutions.

## Results

### Lateral flagella improve chemotactic performance in hydrogels and polymer solutions

We employed a recently developed multiscale chemotaxis assay that 3D-tracks bacteria navigating a linear chemical gradient, allowing simultaneous access to both the population-level chemotactic performance as well as the underlying individual 3D behavioral mechanisms^34^ (Fig. 1b). In order to compare bacterial populations with or without lateral flagella in addition to the polar flagellum, we induced the production of lateral flagella in wildtype *V. alginolyticus* 138-2 by growing it in the presence of the high-molecular weight polymer PVP K90, yielding the “polar + lateral” (PL) phenotype (Fig. 1a). A “polar-only” (P) phenotype is produced by growing mutant YM4^22^ deficient for lateral flagella under the same conditions. We provide a phenotypic assessment in SI Figure 1.

**Fig. 1:**
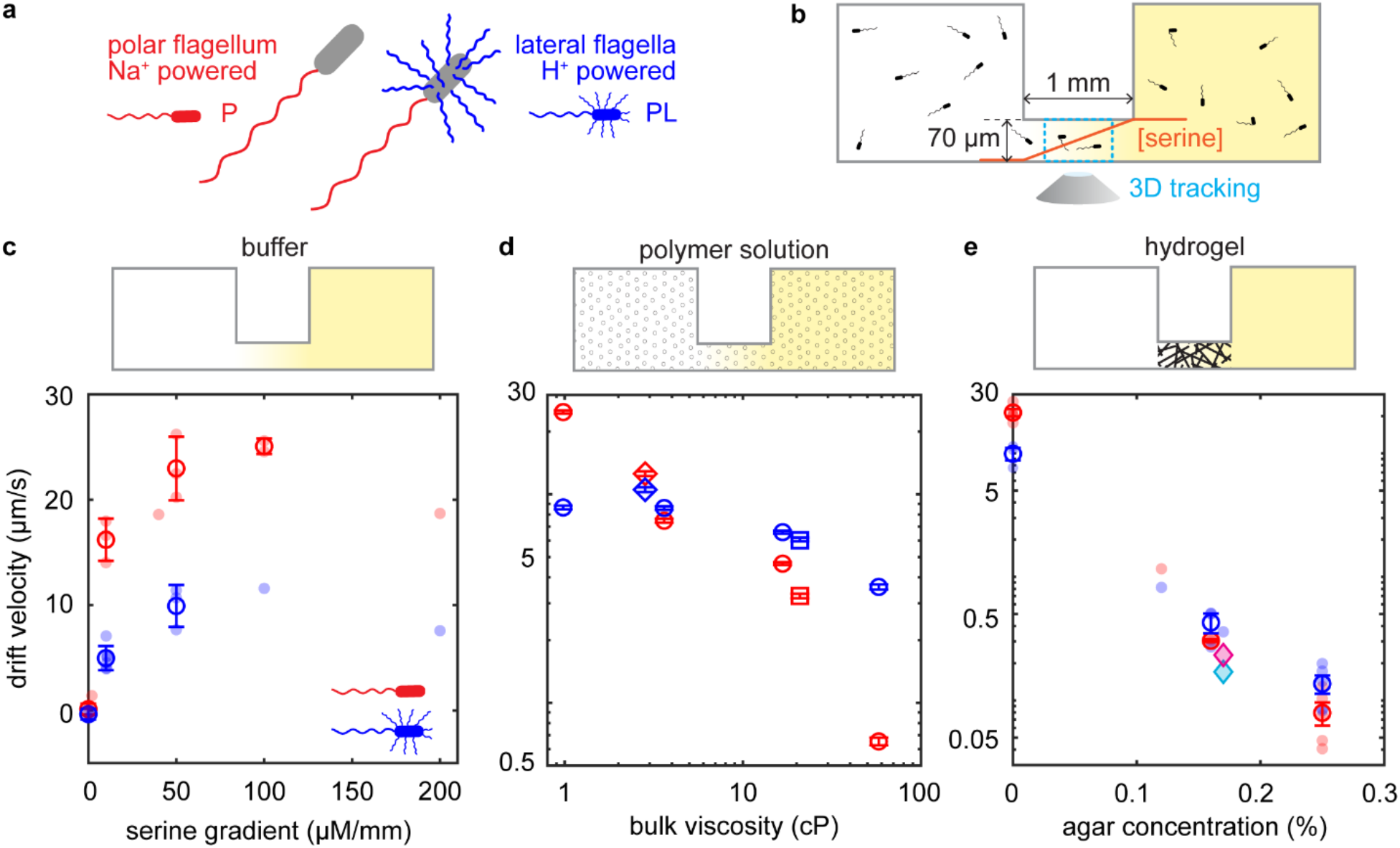
Lateral flagella decrease the chemotactic drift speed in liquid buffer but increase it in complex environments. a) Cartoons of the two flagellar phenotypes and their symbols used later. The polar-flagellum-only phenotype (P, red miniature) is motile by a single Na^+^-driven polar flagellum, while the polar-and-lateral-flagella phenotype (PL, blue miniature) has both a polar flagellum and H^+^-driven lateral flagella. b) Schematic of the multiscale chemotaxis assay. Bacteria are tracked in the middle of a linear serine gradient, established by diffusion in a 1-mm-long, 2-mm-wide, 70-μm-high channel connecting two reservoirs with equal bacterial concentrations, but with or without L-serine. A typical experiment in buffer yields thousands of individual trajectories in minutes. c-e) Schematic of the chemotaxis assays (top) performed in buffer (c), polymer solutions (d), and agar hydrogels (e), and the resulting chemotactic drift velocity of the two flagellar phenotypes (bottom; red: PL, blue, P) as a function of the L-serine gradient in buffer (c) or the complex medium concentration in an L-serine gradient of 50 μM/mm (d,e). Pale symbols in (c,d) represent individual biological replicates. Open circles represent averages between at least two (c), one (d) or three (e) biological replicates. Error bars represent standard deviations between these replicates (c,e) or standard errors of the mean estimated by jackknife resampling a single dataset into subsets consisting of 150 trajectories each (d). In panel (d), the effective bulk viscosity of PVP solutions is shown as a proxy of the concentration. A viscosity of 1 cP approximately corresponds to buffer without PVP. Different symbols represent different biological replicates. In panel (e), cyan and red diamonds represent the wildtype and the Pof-only mutant, respectively, grown without added PVP so as to not induce lateral flagella expression in the wildtype.

We tracked both phenotypes in gradients of L-serine ranging from 0 μM/mm up to 200 μM/mm in TMN buffer (Methods). A typical experiment yielded 3,000 to 6,000 individual trajectories of motile bacteria in about 5 minutes of recordings. We directly compute the chemotactic drift velocity of the population, v_d_, as the average of all signed instantaneous velocity components along the gradient direction, averaged across all time points and all individuals. For both phenotypes, the drift velocity increases as the gradient increases up to approximately 50 μM/mm (Fig. 1c). At all serine gradients tested, the P phenotype displays a higher drift velocity compared to the PL phenotype (Fig. 1c). In a serine gradient of 50 μM/mm, the P and PL phenotypes achieve a chemotactic drift of 23 ± 3 μm/s (mean ± standard deviation (SD), n = 3) and 10 ± 2 μm/s (n = 3), respectively.

We expand the use of the multiscale chemotaxis assay from buffers^34^ to polymer solutions and soft agar hydrogels (Methods). We compare the two phenotypes in varying PVP K90 and agar concentrations with a serine gradient of 50 μM/mm (Fig. 1d,e). For both phenotypes, the drift velocity decreases with increasing concentration, but more rapidly for the P than for the PL phenotype. While the P phenotype outperforms the PL phenotype at low concentrations, the chemotactic performance ranking is inverted above a critical concentration: Above approximately 0.15% agar or 1.4 % PVP K90 (3.6 cP), the PL phenotype outperforms the P phenotype with respect to chemotactic drift velocity (see SI Table 1 for data summary). To rule out an impact of strain differences other than flagellation on chemotactic performance, we grew the wildtype strain without added PVP so as to not induce the expression of lateral flagella. The lack of lateral flagella expression reverses the performance benefit compared to the P strain in 0.17% agar (Fig. 1e).

Thus, at the population scale, lateral flagella decrease the drift velocity in buffer but increase it above a certain polymer or hydrogel concentration. Next, we investigated the behavioral mechanisms underlying the observed chemotactic performance.

### Lateral flagella decrease the drift velocity in buffer by lowering the swimming speed

For a serine gradient of 50 μM/mm in buffer, we extracted 24,248 and 22,101 trajectories of motile individuals (i.e., with a minimum average swimming speed of 18 μm/s) of the P and PL phenotypes, respectively, from three biologically independent experiments each (SI Table 1). In trajectories with a minimum duration of 0.8 s, turning events were identified using an automated turn detection procedure (Methods, SI Table 2). For both the P and PL phenotype, we observe a run-reverse-flick motility pattern as previously described for the P phenotype^5^ (Fig. 2 a,b; SI Fig. 2). Forward and backward runs alternate. The transition from forward to backward swimming is accompanied by a reversal (a turn close to 180°) but the reverse transition can instead produce a turn by a much smaller angle, called a flick. While the phenotypes display indistinguishable turning frequencies of 0.60 ± 0.08 Hz (P, mean ± SD between 3 biological replicates) and 0.61 ± 0.05 Hz (PL), differences between the two populations are apparent in the turning angle distributions (SI Fig. 2c,d) and the average swimming speed (Fig. 2d). The average flick angle is higher for the PL than the P phenotype: Turns by an angle less than 150° average 103° ± 35° (PL, mean ± SD) versus 90° ± 38° (P). Higher flick angles in the presence of lateral flagella have previously been observed for *Shewanella putrefaciens*^24^ and are consistent with the presence of lateral flagella decreasing reorientation during the flick due to increased drag^24,35^.

**Fig. 2.**
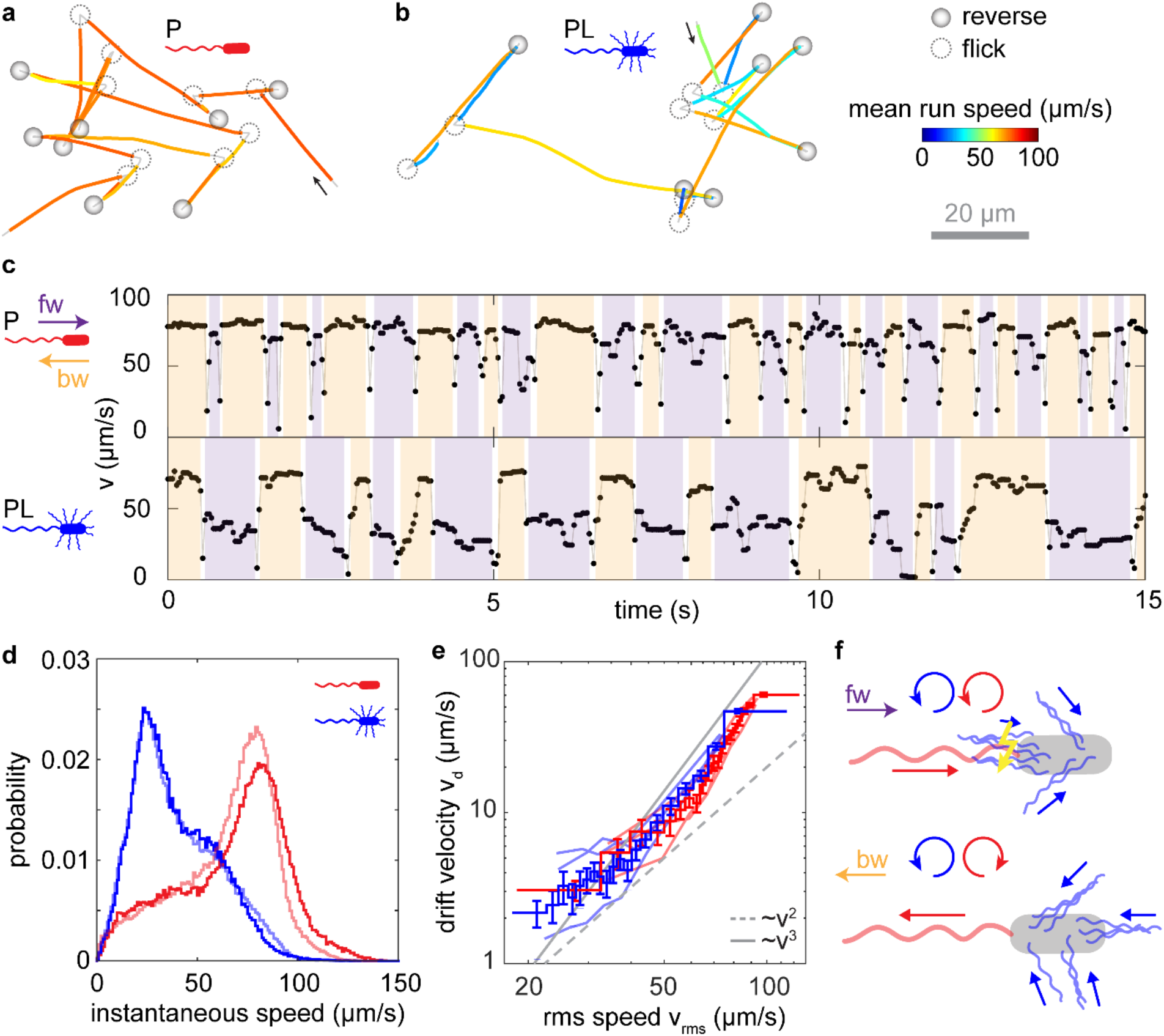
Effect of lateral flagella on motility and chemotaxis in buffer. a-b) Example trajectories for the P (a) and PL (b) phenotypes swimming in buffer, with average swimming speeds similar to the respective population average (see SI Table 2). Run segments are color-coded by their average speed for visual clarity. Turning events are identified and highlighted. SI Fig. 2a, b show the same trajectories color-coded by instantaneous speed. c) Time series of instantaneous speeds for the trajectories in (a) and (b), respectively, where forward (purple) and backward (orange) runs are identified. d) Histograms of instantaneous speed for the motile population of each phenotype (P: red, PL: blue) measured in chemotaxis chambers in a 50 μM/mm serine gradient (light shades) and in motility chambers without gradients. e) Average drift velocity computed for sets of trajectories binned by root-mean-square (rms) speed for the P (red) and PL (blue) phenotypes. Data from individual biological replicates are displayed in light shades, while the stairs with error bars are computed on the pooled data of each phenotype. Trajectories of at least 0.8 s are binned by individual rms speed (8 bins within replicates, 20 bins for the pooled data)) such that each bins contains trajectories of similar cumulative duration, across which rms swimming speed and drift velocity are computed. The error bars reflect the 95% confidence intervals, as estimated by a jackknife resampling procedure consisting of dividing the quantile’s data into subsets of 50 trajectories and computing the standard error of the mean on drift obtained for different subsets. f) Schematic of proposed arrangement and thrust contributions of flagella during forward (top) and backward (bottom) swimming. During forward swimming, lateral flagella interfere with the rotation of the polar flagellum.

The P phenotype displays a higher average swimming speed than the PL, at 65.0 μm/s versus 39.6 μm/s, respectively (Fig. 2d). The decreased average swimming speed of the PL phenotype results from a decreased run speed and affects forward runs more severely than backward runs (Fig. 2c; SI Fig. 2). While the polar flagellum can either push or pull the cell, depending on its rotation direction, the peritrichous lateral flagella are thought to rotate only unidirectionally, pushing the cell^32^. Polar and lateral flagella differ substantially in helical period with values of 1.5 μm and 0.9 μm^36^, respectively, and therefore likely cannot bundle together. The lateral flagella are thus expected to align roughly opposite, respectively parallel, to the polar flagellum during backward, respectively forward, swimming (Fig. 2f). In *S. putrefaciens*, the lateral flagella were observed to maintain typical angles of 30-50° with respect to the trailing end of the cell and to occasionally overlap with the polar flagellum during forward swimming^37^.

We therefore interpret the decreased swimming speed in the presence of lateral flagella in buffer as a combination of two effects: (i) In buffer, lateral flagella may contribute more hydrodynamic drag than thrust, thus decreasing both the forward and the backward swimming speed; and (ii) lateral flagella may interfere with the polar flagellum’s rotation when both are pushing the cell, thus lowering forward swimming speeds more severely than backward speeds (Fig. 2f).

The chemotactic drift velocity is expected to depend not only on swimming speed and turning frequency, but also on turning angles^38,39^, raising the question whether swimming speed differences alone account for the differences in chemotactic performance observed between the P and PL phenotypes. We thus determine chemotactic drift as a function of individual swimming speed for both populations (Fig. 2e). For both phenotypes, the drift velocity increases more steeply with swimming speed than the theoretically predicted quadratic dependence^38^, similar to previous observations in *E. coli*^40^, but in contrast to indirectly inferred drift velocities previously reported for the P phenotype of *V. alginolyticus*^41^. The phenotypes display very similar chemotactic drift at the same swimming speed.

We conclude that the decreased chemotactic drift of the PL phenotype in buffer can largely be attributed to its reduced swimming speed compared to the P population, and that the observed difference in turning angles is of comparatively minor importance for the chemotactic performance in buffer.

### Lateral flagella alter swimming behavior in polymer solutions

Given that, in the linear, high-molecular-weight polymer PVP K90, the PL phenotype outperforms the P phenotype in chemotactic drift speed above a certain macroscopic viscosity, we investigated whether similar effects exist for other polymer solutions at similar viscosities. We quantified the chemotactic drift of both phenotypes in solutions of the lower-molecular-weight PVP K60 as well as in Ficoll 400, a highly branched polysaccharide polymer with a molecular weight similar to the linear PVP K90. We found that the P phenotype maintained a similar or higher chemotactic drift than the PL phenotype at all concentrations of those polymers (Fig. 3a), even when reaching similarly low absolute values as previously observed in PVP K90. PVP K90 is thus the only polymer in which we observe a substantial performance advantage for the PL over the P phenotype.

**Fig. 3:**
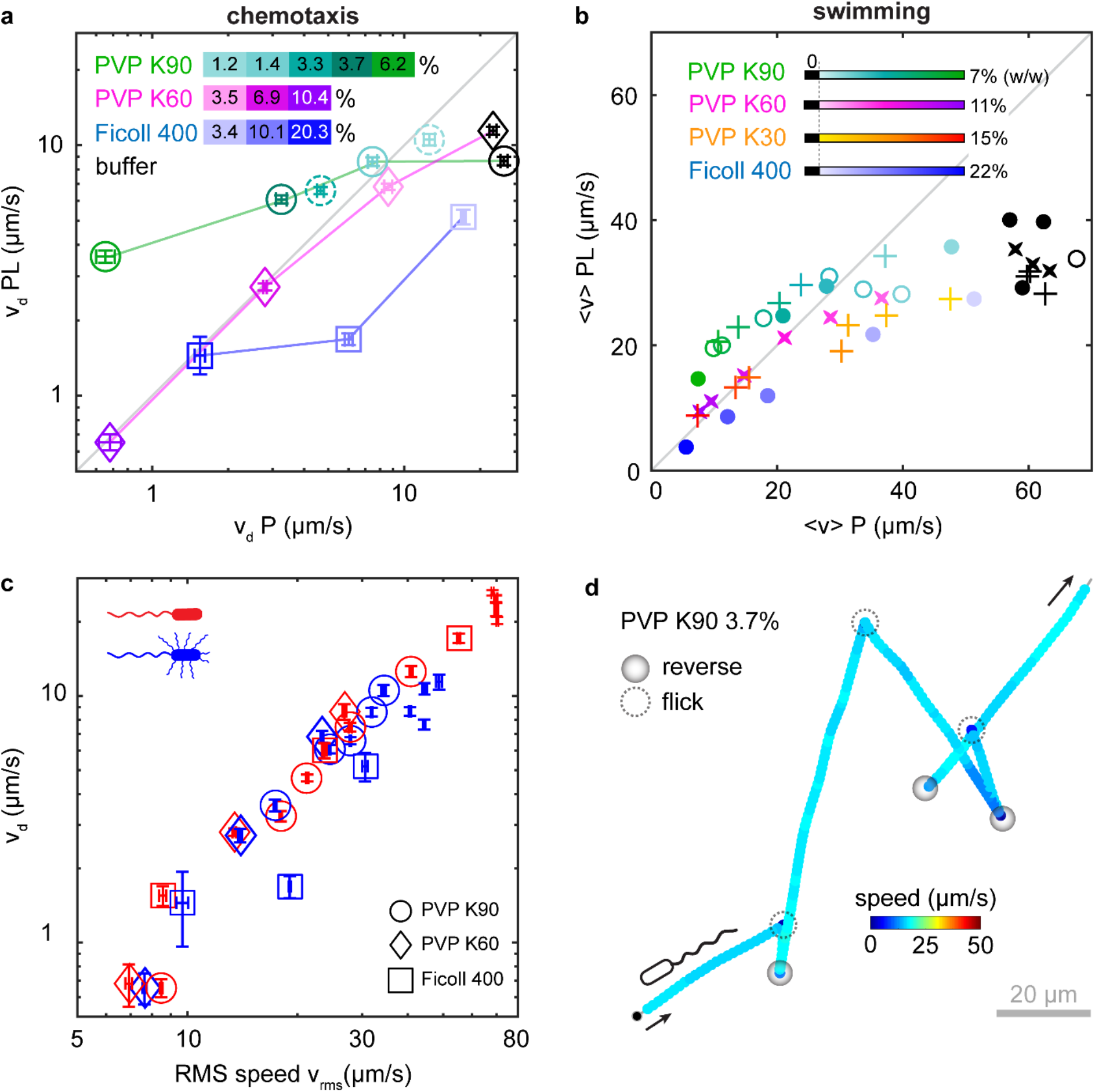
Effect of lateral flagella on motility and chemotaxis in polymer solutions. a) PL drift velocity versus P drift velocity in a 50 μM/mm serine gradient, for the indicated range of concentrations of PVP K90 (circles), PVP K60 (diamonds), and Ficoll 400 (squares). Each point represents a pair of drift measurements for both phenotypes, acquired on the same day in the same media, with error bars estimated by a jackknife resampling procedure consisting of dividing the data into subsets of 150 trajectories and computing the standard error of the mean drift obtained for different subsets. The grey line marks identical performance of both phenotypes. Experiments performed on the same pair of cultures are connected by lines. b) PL average swimming speed versus P average swimming speed at the indicated polymer solutions and concentrations measured in the middle of approximately 300-μm-high motility chambers without chemical gradients. Black symbols represent values observed in buffer solution. Marker shapes indicate biological replicates (the same pair of P and PL cultures). c) Drift velocity given rms population swimming speed for the P (red) and PL (blue) phenotype at a range of concentrations of PVP K90 (circles), PVP K60 (diamonds), Ficoll (squares) or buffer (no symbol). Error bars represent 95% confidence intervals. d) Example trajectory for the P phenotype in 3.7 % PVP K90 (21 cP), displaying run-reverse-flick motility, manually annotated. The trajectory exhibits an average speed matching the population-averaged speed of 17 μm/s and an average flick angle of 123°.

In buffer, swimming speed emerged as the main determinant of chemotactic performance. Previous work found that the PL displays a higher swimming speed than the P phenotype in PVP K90 solutions above a macroscopic viscosity of 3-4 cP^23^. Here, we find that the population-averaged swimming speed and chemotactic performance both exhibit an inversion in the ranking of phenotypes at a similar viscosity of 3.6 cP at 1.4 % PVP K90 (Fig. 1d and SI Fig. 3). At the highest PVP K90 concentrations tested (6.2 %, 58 cP), the PL phenotype swims more than twice as fast as the P phenotype. For PVP K60 and Ficoll 400, there also appears to be a small advantage in swimming speed for the PL over the P phenotype at the highest concentrations tested, but the differences of 10% and 12%, respectively, are comparable in magnitude to the biological variability of 9% (SD/mean across three biological replicates) we observed in buffer. The marginally higher swimming speeds of the PL phenotype compared to the P phenotype do not yield a higher chemotactic drift here, suggesting that factors other than swimming speed may still contribute under these conditions. PVP K90 is thus the only polymer for which we observe a substantial swimming speed and chemotactic advantage for the PL compared to the P phenotype.

In order to investigate the dependence of motility behavior on concentration in polymer solutions in more depth, we performed 3D tracking of both phenotypes in the absence of gradients in approximately-300-μm-high sample chambers, far from surfaces, for a broader range of polymers and concentrations (Fig. 3b, SI Fig. 4a). These data also demonstrate that lateral flagella substantially increase the average swimming speed above a critical concentration of PVP K90, but only yield a minor increase in lower-molecular-weight PVPs, and none in Ficoll 400 (Fig. 3b, SI Fig. 3a). The critical concentration of approximately 2% PVP K90 (6 cP) observed here is a little higher than measured in the chemotaxis assay, perhaps as a result of biological variability between experiments. These observations are also consistent with the notion that differences in the chemotactic performance might be driven by differences in swimming speed.

We also quantified other statistical descriptors of the motility patterns, although the lower swimming speeds and altered refractive indices at high concentrations entail an increased impact of localization errors on the fidelity of the turn detection procedure. Both phenotypes still perform run-reverse-flick motility at all concentrations tested (Fig. 3d, SI Fig. 4e,f). Both phenotypes also display a decrease in turning frequency with decreasing swimming speed (SI Table 2), similar to prior findings on the P phenotype when the swimming speed was modulated via the sodium concentration in the medium^41^. The motility patterns of the P and PL phenotypes become more similar at high concentrations of PVP K90 (SI Fig 4d-f). While the PL phenotype displayed slower forward than backward speeds in buffer, these speeds are similar for both phenotypes at high concentrations of PVP K90 (SI Fig. 4d). In addition, the flick angles become larger for both and approach reversals in magnitude (SI Fig. 4e,f). Both phenotypes thus display a motility pattern closer to “run-reverse” than “run-reverse-flick” at high concentrations of PVP K90. We thus conclude that the higher chemotactic performance of the PL over the P phenotype does not result from differences in turning angles.

More broadly, we can constrain contributions from motility parameters other than swimming speed by comparing P and PL bacteria of similar swimming speeds (S.I. Fig. 4a-c). At the same swimming speed, the chemotactic drift of P bacteria is either higher or similar to the chemotactic drift of the PL bacteria, demonstrating that the higher chemotactic performance of the PL compared to the P phenotype at high concentrations of PVP K90 results from its higher swimming speed. Population averages of chemotactic performance plotted against swimming speed for all polymer conditions tested reveal a global, increasing trend (Fig. 3c), also indicating that swimming speed is the main determinant of chemotactic performance under these conditions.

In order to increase the swimming speed at high concentrations of PVP K90, the thrust from the lateral flagella must outweigh the additional drag they cause, in contrast to the situation in buffer, where we observed a decreased swimming speed of the PL compared to the P phenotype. Interestingly, Martinez et al.^42^ reported that *Escherichia coli* motility reveals non-Newtonian properties in PVP K90 solutions, but not in PVP K30 and Ficoll 400. They proposed that, due to shear thinning of the polymer solution driven by flagellar rotation, the flagella effectively experience a lower viscosity than the cell body, leading to increased swimming speeds. Similar non-Newtonian effects might contribute to increasing the thrust produced by *V. alginolyticus* lateral flagella in high concentrations of PVP K90 so as to outweigh the drag they cause, resulting in the PL phenotype displaying greater swimming speeds than the P phenotype. We, however, also observed small swimming speed advantages of the PL over the P phenotype in PVP K60 and K30, suggesting that additional factors beyond non-Newtonian behavior might contribute, for example different responses of the polar and flagellar motors to increased load^43,44^. Alternatively, the higher flagellar rotation rates of *V. alginolyticus* compared to *E. coli* might probe a different regime of shear rates susceptible to non-Newtonian effects even in lower-molecular-weight PVP.

### Impact of lateral flagella on swimming behavior in a hydrogel environment

At agar concentrations above 0.12%, the PL phenotypes consistently outperformed the P phenotype in chemotactic drift, both on average across all replicate experiments (Fig. 1e) as well as within each pair of experiments within a replicates, except for one where they are within error of each other (SI Fig. 5e). When the lateral flagellar motors were selectively stalled by adding the ionophore CCCP (carbonyl cyanide m-chlorophenyl hydrazine, see Methods for details) which removes the proton gradient driving lateral flagella rotation (SI Fig. 5a,b), the PL population did not outperform the P population anymore (SI Fig. 5c,d). Lateral flagella must therefore play an active role in navigation in soft agar gels.

In agreement with previous observations for *E. coli*^45^, we find that *V. alginolyticus* trajectories in soft agar show drastically different features compared to those in buffer or viscous polymer solutions. Between swimming phases at speeds up to those observed in buffer, the trajectories display typically several-second long “stall” phases of very low speed where the bacteria appear to be trapped (Fig. 4a). We therefore established a new trajectory analysis approach, based on stall and swim phase detection instead of turning event detection (Fig. 4b, Methods). We find that lateral flagella do not increase the speed during the swim phases: the P phenotype is swimming faster than the PL phenotype at all soft agar concentrations tested (Fig. 4c). When the drift velocity is computed based only on the swim phases, the PL phenotype never outperforms the P phenotype (Fig. 4d). As the stall phases appear to be critical for the performance inversion between the phenotypes at increasing agar concentration, we investigated their statistics in more depth.

**Fig. 4.**
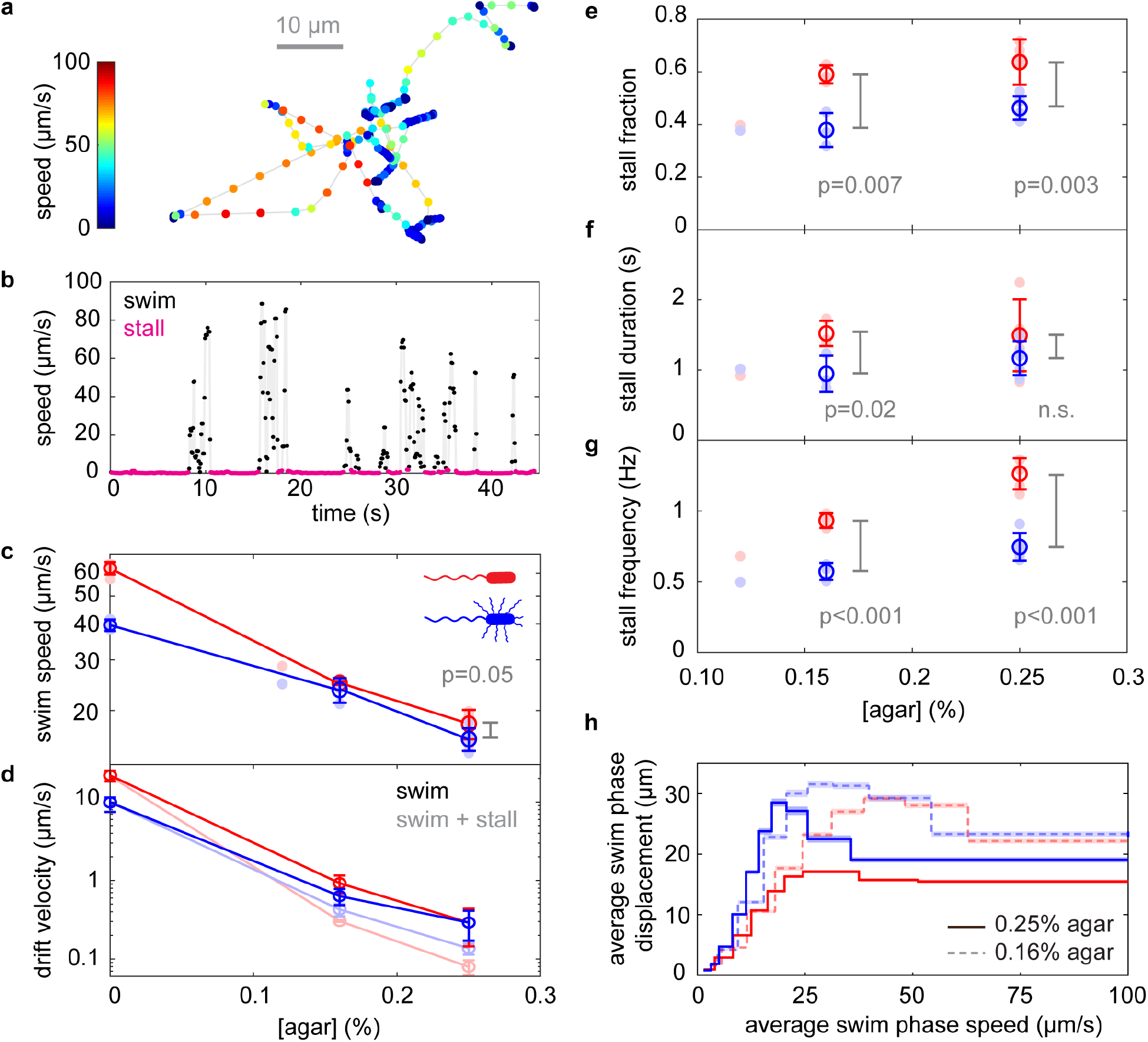
Swim-stall motility in agar. a) A 44.6-s long example trajectory for the P phenotype of with an average speed of 8.6 μm/s in 0.16% agar. b) Time trace of instantaneous speed for the same trajectory. Swim (black) and stall phases (magenta) begin when the bacterium’s instantaneous speed is above, respectively below, 2 μm/s for at least 0.2 seconds. c) Average speed during swim phases plotted against agar concentration for the P (red) and PL (blue) phenotype. The P phenotype swims faster than the PL phenotype during the swimming phases. The lines connect averages between at least three biological replicates, with error bars reflecting SD between biological replicates. d) Drift velocity computed across all entire trajectories (solid) or only during swim phases (faded). e-g) Swim-stall phase analysis for both phenotypes. e) Fraction of time spent stalling, f) average durations of stall phases and g) frequency of stalling when swimming for both phenotypes, plotted against agar concentration. P-values are shown for one-sided t-tests between biological replicates. Error bars reflect SD between biological replicates. f) Average swim phase displacement given the average swim phase speed, in agar 0.16% (dotted lines) and 0.25% (solid lines). For each phenotype and agar concentration, all swim phases from all biological replicates are gathered and distributed equally in number across 10 bins. Across each bin, we average swim phase displacements as the distance between the first and last position in each swim phase. The shading reflects 95% confidence intervals computed as 1.96 times the standard error on the mean. All bins are represented but the fastest bin is graphically cut at 100 μm/s for visual clarity.

We find that the fraction of time spent stalling is 1.4-times higher for the P than the PL phenotype at 0.25% agar (Fig. 4e), raising the question of whether the PL phenotype is stalled less frequently or for shorter durations than the P phenotype. While we observed increased stall durations in the P compared to the PL phenotype above 0.12 % agar, the difference was statistically significant only at 0.16% and not at 0.25% agar (Fig. 4f). On the other hand, the stall frequency, defined as the rate at which swimming bacteria stall, is significantly different between the two populations: on average, lateral flagella decrease the chance of stalling by a factor 1.7 ± 0.2 (mean ± SD) in 0.25 % agar (Fig. 4g). Thus, both the duration and the temporal frequency of stalls are decreased in the presence of lateral flagella.

The stall events likely reflect trapping of bacteria in pores of the agar hydrogel network^45^. Given the lower average swim phase speed of the PL compared to the P population (Fig. 4c), the question arises whether the lower stall frequency of the PL population only reflects the longer time period required to reach the same trap location at a lower speed. Therefore, we quantified the average spatial displacements during swim phases, corresponding to a bacterial “mean free path”, as a function of average swim phase speed for both phenotypes at two agar concentrations (Fig. 4h). At both agar concentrations, the PL phenotype displays longer or similar mean free paths than the P phenotype for similar swim phase speeds. For the fastest 20% of the swim phases, the mean free path of the PL population approaches values similar to those of the P population, perhaps reflecting heterogeneity in lateral flagella expression in the PL population. Individuals with fewer lateral flagella are expected to swim faster and display behavior closer to that of the P population.

## Discussion

In nature, motile bacteria traverse a wide range of environments with complex mechanical properties. The habitats of bacteria that interact with hosts, e.g, as pathogens, symbionts, or commensals, are rich in organic materials whose mechanical properties are often dominated by large polymers. Hosts shield themselves from pathogens using hydrogel networks such as the mucus lining the epithelial surfaces in the respiratory, vaginal, and digestive tracts, or fibrinogen clots sealing wounds. Mucins, the large glycoproteins that form the building blocks of mucus, are not only found in gels, but also in solution, for example in saliva. Solutions and hydrogels of large polymers thus represent two major classes of mechanical environments that motile bacteria encounter in nature. *V. alginolyticus*, like *V. parahaemolyticus*, can infect the digestive tract as well as wounds.

Previous work^11,46^ largely viewed the lateral flagella of *Vibrio* species as a physiological adaptation to surface motility because mutants deficient in lateral flagella do not exhibit swarming motility on the surface of semi-soft agar plates^22^. *Pseudomonas aeruginosa*, however, swarms with solely a polar flagellum^47^, thus lateral or peritrichous flagella are not per se required for swarming. More recent work in *S. putrefaciens* revealed that lateral flagella can also enhance random swimming motility in buffer^24^. Both *V. alginolyticus* and *V. parahaemolyticus* produce lateral flagella under a wide range of environmental conditions, including in polymer solutions and agar gels^12,22^.

Here, we show that lateral flagella enhance chemotactic performance in both these two classes of environments, though by two radically different mechanisms. In the non-Newtonian solutions of PVP K90, lateral flagella improve chemotactic performance by increasing the swimming speed. In soft agar hydrogels, by contrast, lateral flagella do not increase the swimming speed, but increase the mean free path between trapping events, thereby enabling a greater overall chemotactic performance. Figure 5 presents a schematic summary of these behavioral mechanisms.

**Fig. 5:**
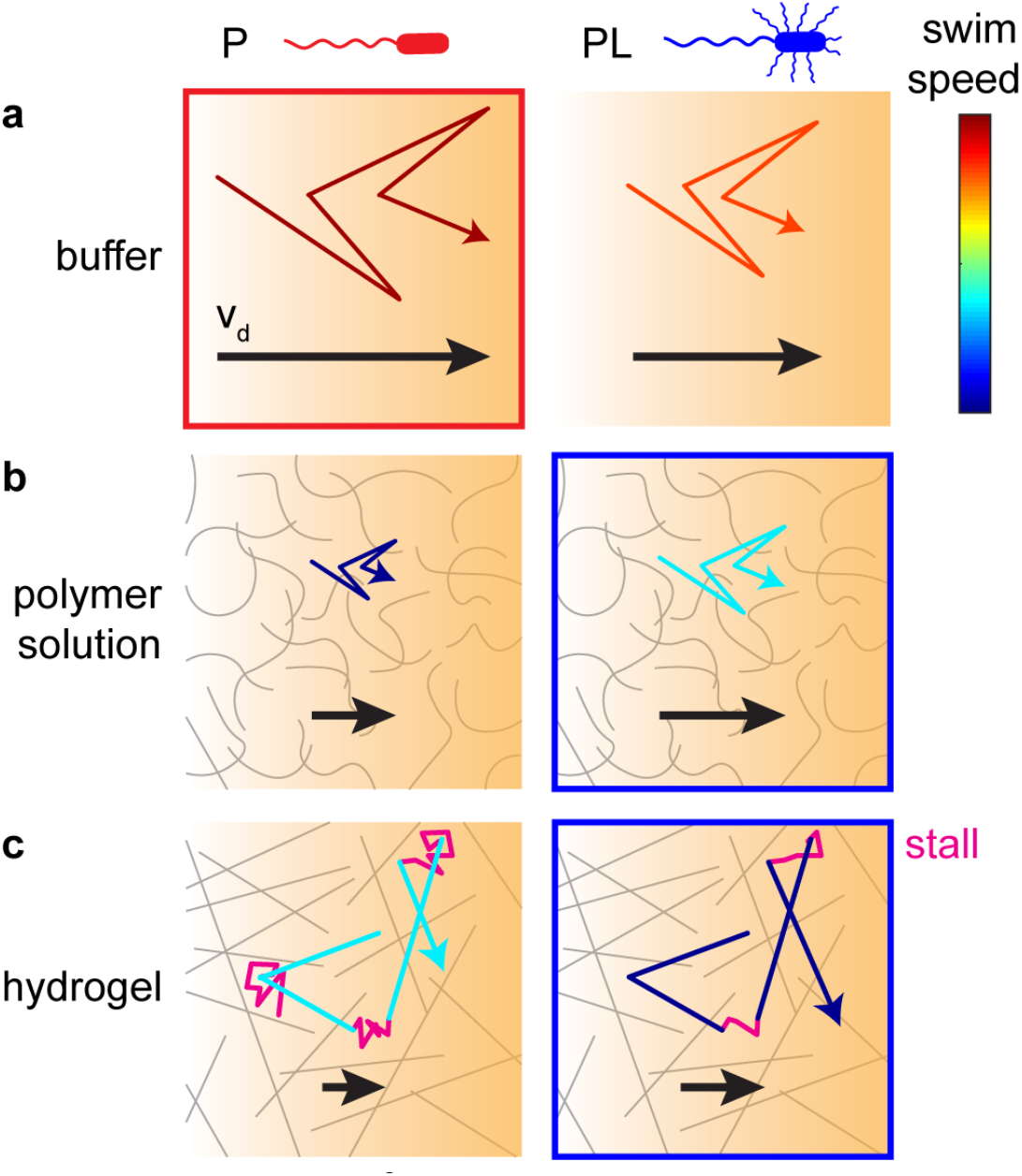
Schematic of behavioral mechanisms that drive chemotactic performance in different environments. a) In buffer, lateral flagella decrease chemotactic performance by decreasing the swimming speed. b) In polymer solutions like PVP K90, lateral flagella increase chemotactic performance by increasing the swimming speed. c) In hydrogels, lateral flagella increase chemotactic performance by decreasing the fraction of time spent stalling. Frames highlight the phenotype enabling the higher chemotactic performance for each environment.

From an evolutionary point of view, our findings seem natural: The expressed phenotype appears to correspond to the phenotype enabling the highest chemotactic performance in a given environment. Lateral flagella, however, do not only mediate benefits but also incur a substantial growth cost: At the concentration of 7% PVP K90 we used to express the PL phenotype in the wildtype strain, this strain displays an approximately 30% longer doubling time than the Pof-only mutant used for the P phenotype under the same conditions (SI Fig. 1f). The ratio of their doubling times increases monotonically with increasing PVP concentration in the growth medium, suggesting that the phenotypic space spanned by the dual flagellar systems may not be as binary as has been assumed but may instead offer a continuum of gradations. In line with this hypothesis, previous work in *V. parahaemolyticus* has shown that the induction rate of lateral flagella varies continuously with environmental conditions^13^.

Our chemotaxis assay measures only the benefits of lateral flagella and the growth measurements only the cost. How do the bacteria navigate these two conflicting demands? A commonly used assay that integrates the demands for rapid growth and efficient chemotaxis is colony expansion in soft agar plates^48,49^. We measured the colony expansion rate in a range of soft agar concentrations for both strains. We find that the wildtype strain expands faster than the Pof-only mutant at the commonly used agar concentrations of 0.2-0.3% agar, but the Pof-only mutant actually outperforms the wildtype at agar concentrations below 0.16% (SI Fig. 1e). Intriguingly, this value is a little higher than the agar concentration between 0.12% and 0.16% at which we observed an inversion in purely chemotactic performance between the P and PL phenotypes (Fig. 1d). The deviation might indicate that, at the lower agar concentration, the chemotactic advantage mediated by lateral flagella does not yet outweigh their growth cost. In addition, the fact that the wildtype is outperformed by the Pof-only mutant at low agar concentrations indicates that the wildtype’s expression of lateral flagella might not necessarily be fully optimized towards the currently experienced conditions. Even when grown in medium without any added viscous agents, the wildtype strain expresses low levels of lateral flagella (SI Fig 1d) and displays a higher doubling time than the Pof-only mutant (SI Fig. 1f). It is unclear at present whether this low expression level is uniform across individuals or reflects phenotypic heterogeneity that enables bet hedging in the face of unpredictable environmental change. Another open question is whether bacteria distinguish between the non-Newtonian polymer solutions in which lateral flagella offer an advantage and Newtonian polymer solutions where they do not. Mucus is also non-Newtonian and exhibits shear thinning^50^, suggesting that these properties might even be so common in nature as to render the distinction unnecessary for the bacteria.

Our findings represent first steps towards a more systematic exploration of the rich landscape of costs and benefits of flagellar architectures and the navigation strategies they enable in natural environments.

## Methods

### Bacterial strains and growth conditions

Bacterial strains are listed in SI Table 3. Overnight cultures were inoculated in 2 ml MB (Difco Marine Broth 2216, passed through a 0.2 μm filter to remove precipitate) from individual colonies grown on 1.5-2% agar MB plates streaked from frozen glycerol stocks, and grown to saturation at 30°C, 250 rpm. Day cultures were inoculated with the overnight cultures at 1:200 dilution in 10 ml MB with or without 7% PVP K90 and grown at 22°C, 200 rpm, until they reached an optical density (OD) at 600 nm between 0.34 and 0.40, except when noted otherwise. Volumes of 1 ml of bacterial culture without PVP K90 were washed once by centrifugation in 1.5 ml microcentrifuge tubes (7 min at 2,000 rcf), followed by gentle resuspension in 1 ml of motility medium TMN (50 mM Tris-HCl, 300 mM NaCl, 5 mM MgCl_2_, 5 mM glucose, pH 7.5). For day cultures with PVP K90, 0.5 ml of bacterial culture was combined with 0.5 ml TMN and slowly mixed by agitation to decrease the solution’s viscosity, then washed by centrifugation in 1.5 ml microcentrifuge tubes (20-28 min at 2,000-2,600 rcf), followed by gentle resuspension in 1 ml of TMN. They were diluted to a target OD of approximately 0.005 for chemotaxis experiments in buffer or polymer solutions, or 0.001 or less for chemotaxis experiments in agar, either in TMN for chemotaxis experiment in buffer or agar or in TMN with the polymer concentration, with or without chemoattractant, for injection into the chemotaxis device.

### Viscous polymer solutions

PVP K90 & K30 (powder, Sigma-Aldrich) prepared respectively as 14% and 30% solution in ultrapure water, PVP K60 (45% solution in water, Sigma-Aldrich) and Ficoll Type 400 (20% solution in water, Sigma-Aldrich) stocks were all dialyzed. 30 to 50 ml of the stock solution was placed in 70 ml dialysis cassette with a 10kDa cut off (Slide-A-Lyzer G2 cassettes, Thermo Scientific), then let at least 15 days in 4 or 5 L beaker filled with ultrapure-water and slow agitation, with at least daily water change. To obtain satisfactory concentrations of stock solutions of dialyzed polymers, they were then reconcentrated after dialysis, using centrifugal filter units with a 10kDa cutoff (Centriprep by MilliporeSigma or Macrosep by Pall Laboratory). The final stock solution concentration was determined by weighting at least three different 1.5 ml tubes with 100-200 μL of the solution, before and after drying in vacuum centrifuge at 80°C for 8 hours. These stocks were used in experiments within 15 days of dialysis, except for the PVP K90 used in our growth medium (MB + 7% PVP K90). We used a previously established relationship between PVP K90 concentration and bulk viscosity^51^.

### Chemotaxis assay preparation in buffer or polymer solution

The chemotaxis assays were performed using a high-throughput chemotaxis assay^34^ where bacteria are tracked in 3D in a controlled chemical gradient established in a commercially available microfluidic device (μ-slide Chemotaxis, IBIDI), consisting of two reservoirs of different chemical concentrations connected by a small channel in which a linear gradient is formed. The device’s reservoirs were filled with the two bacterial solutions (with and without chemoattractant) following a modified version of the manufacturer’s “Fast Method” protocol. First, the entire device was overfilled with buffer free of chemoattractant or bacteria through the filling ports, and then the central channel’s ports were closed with plugs. 65 μl was removed from one reservoir, replaced by 65 μl of chemoattractant-free bacterial solution, and then this reservoir’s ports were closed. Finally, all liquid was removed from the other reservoir and replaced with bacterial solution containing chemoattractant. Key to reproducible gradients is to not overfill this reservoir to avoid liquid flow in the central channel when the last two ports are closed. For experiments in viscous media, the same protocol was followed except for the additional presence of the viscous agent at identical concentration in all solutions. Each biological replicate compares flagellar phenotypes using identical polymer solutions to limit technical variability.

### Chemotaxis assay preparation in agar

For chemotaxis assays performed in agar, the central channel of the microfluidic device was filled in advance with molten agar following a modified version of the manufacturer’s Application Note 17, “Seeding Cells in a Gel Matrix” (https://ibidi.com/img/cms/support/AN/AN17_Chemotaxis2Dand3D.pdf), without any bacteria inside the agar gel. First, with all empty reservoirs closed, the central channel was filled with molten agar: 10 μl of hot soft agar at the desired concentration in TMN was applied at the opening of the bottom port, then air and agar were aspirated slowly from the top port until agar completely filled the central channel and its two ports. The ports were closed immediately, and the device left at room temperature for 45 to 60 minutes to let the agar solidify. The reservoirs’ ports were then carefully opened and filled with 65 μl of buffer each. The devices were stored at room temperature in a closed petri dish containing a moist tissue to limit evaporation and used for chemotaxis experiments within 4 hours. For filling the device, each reservoir is successively emptied, filled with bacterial solution with or without chemoattractant, and then closed, with caution to not overfill the reservoirs. Biological replicates comparing flagellar phenotypes at a given agar concentration were prepared in parallel to limit the influence of technical variability in agar gel properties.

### Motility experiments

After washing and resuspension in TMN as described above, cells were diluted by 1:50 to 1:100 in TMN or TMN with dialyzed polymer solution, incubated at room temperature for at least 15 minutes, to allow adaptation to the medium. Sample chambers with a height of approximately 300 μm were created by using three layers of parafilm as spacers between a slide and a cover glass^35^, filled, sealed with valap and immediately brought to the microscope for data acquisition.

### Data acquisition

Phase contrast microscopy recordings were obtained at room temperature (~22°C) on a Nikon Ti-E inverted microscope using an sCMOS camera (PCO Edge 5.5) and a 40x objective lens (Nikon CFI S Plan Fluor ELWD 40x ADM Ph2, correction collar set to 1.2 mm to induce spherical aberrations^35^) focused at the center of the channel in all three dimensions. The integration time was 5 ms. The frame rate was at 30 fps for recordings in buffer, 15 fps in agar, or adjusted between 10 to 30 fps depending on the observed speed in polymer solutions. Typical cumulated acquisition time for one condition in one chemotaxis experiment (one point in Figure 1c-e) ranged from 4 to 10 minutes in buffer, 6 to 12 minutes in viscous media, 15 to 20 minutes in agar.

### Data processing and 3D tracking

Video recordings were binned by a factor of 2 × 2 pixels by averaging pixel counts and then subjected to a background correction procedure based on dividing the image by a pixel-wise median computed across a sliding time window of 101 frames, except for data acquired for agar experiments, where a sliding window of 3001 frames was used to ensure that stalled bacteria were not erased.

3D trajectories were extracted from phase-contrast recordings using a high-throughput 3D tracking method based on image similarity of between diffraction rings of bacteria and those in a reference library with known vertical positions^35^. For tracking in agar, individual bacteria reside in the tracking volume for a long time because they spent a large fraction of their time stalled. We found that the previously established 3D tracking algorithm^35^ often terminated trajectories prematurely, presumably due to the presence of background noise from other bacteria, leading to biased stall duration statistics. We improved the 3D tracking algorithm by correcting images for the diffraction rings of already tracked bacteria. Every time a bacterial 3D position is determined, a 128×128-pixel region of interest (ROI), I, centered around the bacterial in-plane (x,y) position is corrected using a same-size ROI, R, from the reference library image corresponding to the bacterium’s vertical (z) position. We then approximate the diffraction pattern of the detected bacteria by scaling the intensity level of the reference image. First, we compute the deviation from the background in an image I as I_dev_ = I/median(I) – 1. We define weights W = R_dev_^2^, and compute a scalar, weighted ratio of deviations from background between image and reference library as d = sum(I_dev_ W / R_dev_)/ sum(W). Capital variable names here are 128×128 matrices, and multiplications and divisions on them are performed elementwise. The factor d can be understood as a scaling factor between the reference image and the bacterial image. We then compute the intensity-adjusted reference image as R_adj_ = d R_dev_ + 1. The final corrected image, I_c_, is then computed as I_c_ = I / R_adj_. A corrected version of all tracked image frames is saved in a temporary folder and continuously updated during tracking. Benchmarking based on tracking *E. coli* swimming in buffer revealed a more than 60% increase in the number of trajectories longer than 5 s compared to the previous algorithm.

Positions were smoothed using 2^nd^ order ADMM-based trend filtering^52^ with regularization parameter λ = 0.1 in agar and λ = 0.6 otherwise. Three-dimensional velocities were computed as forward differences in positions divided by the time interval between frames.

### Run-reverse-flick analysis in buffer

The turning event detection is based on the local rate of angular change, computed from the dot product between the sums of the three consecutive velocity vectors preceding and subsequent to a time point. The threshold for a turn to begin is an α-fold rate relative to the median rate of angular change of the run segments, as determined in three iterations of the procedure. We determined by visual inspection of both phenotypes’ trajectories that a factor α = 6 (P) or α = 8 (PL) gave satisfactory results. A new run begins with at least two time points (at least 0.066 s) under this threshold. The 3D turning angle for a turn beginning at frame i and ending at frame j is computed as the angle between the sum of the instantaneous velocity vectors at frames i-2 and i-1 and the sum of those at frames j and j+1. Backward (CCW rotation) and forward (CW rotation) runs were identified as runs with a turn under 130°, respectively at the end or at the beginning the run, and a turn above 150° at the other end of the run. All trajectories with a minimum speed of 18 um/s and with a minimum duration of 0.8 s were analyzed. From the 15,108 (P) or 15,653 (PL) analyzed trajectories in the chemotaxis triplicates in liquid buffer, 2,611 forward and 2,601 backward runs (P) or 3,870 forward and 3,944 backward runs (PL) were identified out of 10,468 (P) or 20,272 (PL) total detected runs.

### Run-reverse-flick analysis in polymer solutions

The turning event detection and trajectory analysis is identical to that in buffer, except for the factor α adjusted to 8 (PVP K90 3.3%) or 10 (PVPK90 6.2%) and the minimum trajectory duration adjusted to 1 second.

### Stall-swim analysis in soft agar hydrogels

The stall event detection is based on the instantaneous velocity.: If the speed is under 2 μm/s for 0.2 s or more, a stall starts. During a stalling event, if the instantaneous speed exceeds 2 μm/s for 0.2 s or more, the stall ends, and a new swimming phase starts. All trajectories of with a minimum duration of 1 s are analyzed.

### Protonophore CCCP experiment

The protonophore carbonyl cyanide m-chlorophenylhydrazone (CCCP, Sigma-Aldrich) was used to collapse the H^+^ motive force used by the lateral flagella while maintaining the Na^+^ motive force powering the polar flagella, as described in Kawagishi *et. al*.^22^. They detail that *V. alginolyticus* has respiration-coupled H^+^ and Na^+^ pumps, the later only active in an alkaline pH range. At pH 7, the Na^+^ motive force is secondarily generated by the Na^+^/H^+^ antiporter from the H^+^ motive force, therefore addition of CCCP at pH 7 also collapses the Na^+^ motive force. But at pH 8.7, the Na^+^ motive force is generated by the primary Na^+^ pump and therefore does not collapse upon CCCP addition. Fresh solutions of CCCP in TMN, buffered at pH 8.7 with TAPS instead of Tris, were prepared on the day of the experiment by dilution from 92 mM CCCP stock in DMSO into a 100 μM solution and subsequent dilutions. We chose to work at 20 μM CCCP in liquid buffer, as the motility of the Laf-only strain was completely lost, but the P and PL populations were still swimming at speeds comparable to those in the absence of CCCP (S.I. Fig 4 a,b).

### Growth rate measurements

The growth burden imposed by lateral flagella was determined in an automated growth experiment in a MultiskanTM FC microplate photometer (Thermo Scientific, software SkanIt 4.1). The wells of a 96-well microplate were filled with 200 μl of either MB alone (blank wells and border wells) or MB with PVP K90 up to 7 % (w/w). 1 μL of saturated cultures of either wild-type or Pof-only strains were added to the wells, except for the blanks (20 wells). Each strain and viscosity combination was represented by at least 5 wells in one experiment. A BreathEasy^R^ sealing membrane (Sigma-Aldrich) was placed on top of the 96-well microplate to limit evaporation. Then the microplate was placed in the microplate reader and incubated at 26°C with highest continuous shaking. Absorbance at 620 nm was recorded every 5 min for the first 8 to 10 h and then every 15 min for another 3 to 5 h.

### Flagella extraction

For each strain and condition, 10 ml of cell culture were collected at OD_600nm_ between 0.4 and 0.5, washed twice in TMN (1,900 rcf, 20 min) and resuspended in TMN with a volume of 2-2.5 ml to reach a similar final cell concentration (2 ml for an initial OD_600nm_=0.4, 2.5 ml for OD_600nm_=0.5). We then sheared the flagella by repeated passage through two 5-ml syringes connected by 22-inch-long tubing (ID 0.023”, BD INTRAMEDIC, BD Medical), about 30 times. 1 ml of TMN is used to rinse off flagella and cells from the tubing. The solution is centrifugated at 4,000 rcf for 25 minutes to pellet the cell bodies. The supernatant, containing sheared flagella, is collected and placed in a centrifugal filter unit (10 kDa Ultra-4 Amicon, Merk Millipore) topped with PBS to 4 ml, then centrifugated at 1,900 rcf for 30 minutes to concentrate the flagella extract. 3.5 ml of PBS is added again and the sample is centrifugated for an additional hour, for sample desalting. About 30 μL of sample is collected (with careful agitation to collect even flagellar components deposited at the bottom of the sample compartment) and adjusted to a final volume of 45 μL.

### SDS-PAGE

Sodium dodecyl sulfate polyacrylamide gel electrophoresis (SDS-PAGE) was employed to visualize flagellar components. First, the 45-μL freshly-extracted samples were mixed with 15 μL of 4x Laemmli sample buffer (Bio-Rad) for a final concentration of 5 % β-mercaptoethanol. The mixture was denatured at 95°C for 10 min then put on ice and stored in the fridge. Once all samples were gathered, 8 μL of sample or buffer or protein standards solution (Precision Plus Protein Kaleidoscope, Bio-rad) were loaded per well of pre-cast 12 % polyacrylamide gel (10-well Mini-PROTEAN TGX, Bio-rad). Tris-glycine buffer (pH 8.3) was used as the running buffer, and the gel was run at 180 V for about 35 minutes (Mini-PROTEAN electrophoresis cell and PowerPac power supply, Bio-rad). Coomassie brilliant blue (Bio-Safe Coomassie stain, Bio-rad) staining allowed visualization of the proteins in the gel.

### AFM sample preparation and imaging

1 ml of day culture of wild-type strain in MB + 7% PVP K90 at OD_600nm_≈0.35 was washed as described above and resuspended in 0.1 ml TMN. 2 μL of glutaraldehyde was added by gentle mixing and incubated at room temperature for 10 minutes. Then 10 μl of the solution were placed onto a clean coverslip and left for 30 minutes to an hour. The coverslip was then carefully rinsed three times with TMN, then 6 times with ultrapure water, and left to dry at 37°C for three hours. The sample was observed and images acquired with a Park NX12 (Park Systems, software SmartScan, non-contact mode), using a non-contact probe (200AC-NA, OPUS, sold by NanoAndMore USA). The image shown in SI Fig. 1 is 256×256 pixel or 15×15 μm. Analyses of flagellar thickness were performed using the XEI software (version 4.3.0. Build5, Park Systems).

### Colony expansion assay in soft agar plates

MB medium and the relevant amount of agar (0.12-0.27 g per 100 ml) were mixed and autoclaved for 30 minutes. Upon cooling down to 55°C, soft agar plates were prepared by pipetting 25 ml into each 90-mm petri dish, immediately closing their lids, and letting it solidify for an hour. 0.5 μL of the relevant overnight culture (prepared as detailed in the section Bacterial strains and growth conditions above) was then carefully injected in the middle of each plate and let to sit for 10 minutes. Automated time-lapse images of plates were recorded every 10 to 15 minutes for up to 20 hours, using a custom-built dark-field imager with a rotating platform holding 8 petri dishes, placed inside an incubator held at 30°C. Colony expansion rates within the soft agar were extracted from a linear fit to the time series of outer ring radii obtained using a custom MATLAB ring-detection algorithm based on Hough transforms. Data obtained at 0.12% agar results from three biologically independent experiments. Data for higher concentrations reflect one soft agar plate per concentration injected with the same overnight culture for each strain. Comparisons between strains at a given agar concentration are performed for soft agar plates prepared identically and simultaneously.

## Acknowledgments

All strains used were a kind gift of Professor Seiji Kojima (Nagoya University, Japan). We thank Alan Stern, Alan Sliski, and Chris Stokes for help with the automated dark field imager for colony expansion assays. This research was supported by the Rowland Institute at Harvard University.

## Author contributions

K.M.T. and M.G. conceived the research. L.E. and M.G. developed, performed, and analyzed the soft agar plate experiments. J.W.N. improved the 3D tracking algorithm. M.G. performed all other experiments and analysis. M.G. and K.M.T. interpreted the data and wrote the manuscript. All authors commented on the manuscript.

## Data availability statement

All trajectory data will be made available on the Harvard Dataverse (www.dataverse.harvard.edu/dataverse/tautelab).

