## Supplementary Information for "Physiological adaptation in flagellar architecture improves *Vibrio alginolyticus* chemotaxis in complex environments"

#### SI Table 1

| data-set # | medi-um | concen-tration | main Fig. | # of motile trajectories | total motile trajectory duration (s) | motility threshold (μm/s) | average speed (μm/s) | $v_d$ (μm/s) |
|---|---|---|---|---|---|---|---|---|
| 1 (n=3) | TMN | | Fig. 1c<br>Fig. 2<br>SIFig. 2 | 24,248<br>22,101 | 33,980<br>51,855 | 18 | 65.0<br>39.6 | 23.0±3SD<br>9.9±2SD |
| ⁺1a | | | | 10,085<br>6,434 | 14,514<br>15,503 | | 64.0<br>39.6 | 26.2±0.3SE<br>10.7±0.3SE |
| $1b | | | | 11,667<br>11,986 | 15,864<br>29,360 | | 65.9<br>38.8 | 20.2±0.3SE<br>7.6±0.2SE |
| °1c | | | | 2,496<br>3,681 | 3,601<br>6,992 | | 64.9<br>42.6 | 22.5±0.7SE<br>11.4±0.4SE |
| *2a | PVP K90 | 0.98 cP (0%) | Fig. 3a,c<br>SIFig. 3<br>SIFig. 4 | 6,749<br>9,473 | 9,711<br>20,388 | 18 | 64.9<br>36.1 | 24.8±0.4SE<br>9.1±0.1SE |
| 2b | | 2.9 cP (1.2%) | | 4,283<br>3,073 | 10,286<br>8,396 | 12 | 41.1<br>31.93 | 12.6±0.3SE<br>10.5±0.3SE |
| *2c | | 3.6 cP (1.4%) | | 7,286<br>10,077 | 21,094<br>22,854 | 10 | 25.4<br>30.6 | 7.5±0.2SE<br>8.6±0.2SE |
| *2d | | 17 cP (3.3%) | | 13,283<br>15,804 | 50,066<br>42,710 | 8 | 19.3<br>26.8 | 4.6±0.1SE<br>6.6±0.1SE |
| 2e | | 21 cP (3.7%) | | 10,007<br>13,224 | 46,809<br>39,161 | 8 | 16.9<br>23.4 | 3.2±0.1SE<br>6.1±0.1SE |
| *2f | | 58 cP (6.2%) | | 5,646<br>6,687 | 54,043<br>30,698 | 2 | 7.7<br>16.3 | 0.65±0.03SE<br>3.6±0.1SE |
| °3a | PVP K60 | 0% | Fig. 3a,c<br>SIFig. 3<br>SIFig. 4b | 2,496<br>3,681 | 3,601<br>6,992 | 18 | 64.9<br>42.6 | 22.3±0.7SE<br>11.5±0.4SE |
| °3b | | 3.5% | | 2,429<br>4,119 | 8,640<br>15,403 | 11 | 24.6<br>21.8 | 8.7±0.3SE<br>6.9±0.2SE |
| °3c | | 6.9% | | 1,467<br>2,877 | 11,633<br>17,230 | 8 | 12.1<br>12.7 | 2.74±0.05SE<br>2.72±0.09SE |
| °3d | | 10.4% | | 945<br>1,602 | 12,942<br>17,430 | 3 | 6.0<br>6.6 | 0.68±0.07SE<br>0.67±0.05SE |
| ˟4a | Ficoll 400 | 3.4% | Fig. 3a,c<br>SIFig. 3<br>SIFig. 4c | 2,692<br>3,256 | 4,973<br>9,266 | 12 | 50.6<br>27.6 | 17.0±0.4SE<br>5.3±0.3SE |
| ˟4b | | 10.1% | | 3,473<br>7,500 | 14,833<br>27,449 | 5 | 21.3<br>17.3 | 6.0±0.2SE<br>1.68±0.08SE |
| ˟4c | | 20.3% | | 774<br>901 | 9,803<br>8,082 | 2 | 7.7<br>9.1 | 1.54±0.07SE<br>1.46±0.25SE |
| 5a | agar | 0.12% | Fig. 1d<br>Fig. 4<br>SIFig.6e | 15,513<br>17,286 | 113,755<br>91,445 | 0 | 18.1<br>16.1 | 1.16 ±0.03SE<br>0.82±0.04SE |
| 5b (n=3) | | 0.16% (n=3) | | 34,696<br>70,786 | 395,111<br>390,173 | | 9.4±0.7SD<br>15.4±3.4SD | 0.30±0.01SD<br>0.43±0.08SD |
| 5c (n=5) | | 0.25% (n=5) | | 16,693<br>21,380 | 360,148<br>289,571 | | 6.4±2.4STD<br>8.9±1.3STD | 0.07±0.02SD<br>0.14±0.02SD |
| ⁋6a | | 0.17% | Fig. 1d<br>SIFig.5e | 6,077<br>9,319 | 91,991<br>109,636 | | 7.4<br>9.4 | 0.23±0.01SE<br>0.17±0.02SE |
| ⁋6b | | | | -<br>6,907 | -<br>81,545 | | -<br>16.0 | -<br>0.36±0.03SE |
| ‡6c | | 0.20% | SIFig.5 | 926<br>1,517 | 29,314<br>7,200 | | 5.4<br>15.5 | 0.15±0.02SE<br>0.23±0.04SE |
| ‡6d | | | | 1,858<br>2,601 | 66,999<br>59,400 | | 5.4<br>6.7 | 0.11±0.01SE<br>0.06±0.01SE |

SI Table 1: Trajectory datasets obtained in chemotaxis chambers with a 50 μM/mm L-serine gradient for the P (first line, red) and PL (second line, blue) phenotypes. The error is given as SD when computed over averages of independent biological replicates, or as SE when estimated by a jackknife resampling procedure. Superscript markers indicate measurements performed on the same bacterial cultures. The absence of a superscript indicates that the culture was not used for any other dataset displayed in this table. n indicates the number of independent biological replicates. For dataset 6a, both strains were grown in MB without added PVP so as to not induce lateral flagella. Datasets 6 were recorded at pH8.7 instead of pH7.5, and in the absence (6c) and presence (6d) of 20 µM CCCP to disrupt the proton motive force driving lateral flagella rotation.

#### SI Table 2

| data-set (#) | me-dium | concen-tration | figure | # of ana-lyzed trajec-tories | total ana-lyzed trajec-tory dura-tion (s) | ave-rage speed (µm/s) | turning fre-quency ($s^{-1}$) | backward run speed (µm/s) | forward run speed (µm/s) |
|---|---|---|---|---|---|---|---|---|---|
| $^{+\circ\$}$**1** | TMN | | Fig. 1c<br>Fig. 2<br>SI Fig. 2 | 15,108<br>15,653 | 28,801<br>46,502 | 65<br>39 | 0.60±0.08 SD<br>0.61±0.05 SD | 65.0±0.4SE<br>50.5±0.3SE | 56.0±0.4SE<br>41±0.3SE |
| $^{+}$**7** | | | Fig. 2d | 1,558<br>1,184 | 5,796<br>3,215 | 67<br>39 | 1.4<br>0.7 | 77.5±0.4SE<br>54.9.0±0.7SE | 69.0±0.4SE<br>41.3±0.6SE |
| **($^{+}$7)** | | | Fig. 2a,b<br>SI Fig. 2a,b | 1<br>1 | 16.6<br>17.3 | 64<br>39 | 2.0<br>1.2 | 77±1SE<br>60±6SE | 70±1SE<br>33±2SE |
| ***2a** | PVP K90 | 0% | SI Fig. 4d-f | 4,155<br>6,597 | 8,205<br>18,508 | 64.9<br>36.1 | 0.63±0.01SE<br>0.42±0.01SE | - | - |
| ***2d** | | 3.3% | | 10,170<br>12,355 | 46,696<br>40,093 | 19.3<br>26.8 | 0.40±0.01SE<br>0.09±0.01SE | - | - |
| ***2f** | | 6.2% | | 5,038<br>5,790 | 53,431<br>29,801 | 7.7<br>16.3 | 0.41±0.01SE<br>0.10±0.01SE | - | - |

SI Table 2: Statistics of trajectory datasets on which run-turn analysis was performed. Superscript markers are consistent with SI Table 2. Dataset 7 was obtained in motility chambers (Methods) without a gradient, all other datasets were obtained in chemotaxis chambers with a 50 µM/mm L-serine gradient. Asterisks denote biological replicates consistent with SI Table 1.

#### SI Table 3

| Name | Phenotype | Designation in this study | Refer-ence |
|---|---|---|---|
| 138-2 | Wildtype. Expresses lateral flagella in viscous media. | PL, if grown in viscous medium | 1 |
| YM4 | No lateral flagella | P, Pof-only | 2 |
| YM19 | No polar flagellum | L, Laf-only | |
| YM18 | No flagella | NF | |

SI table 3: Strains used in this study as well as their phenotypes and designations. YM4, YM19, and YM18 were derived from 138-2.

#### S.I. Figure 1

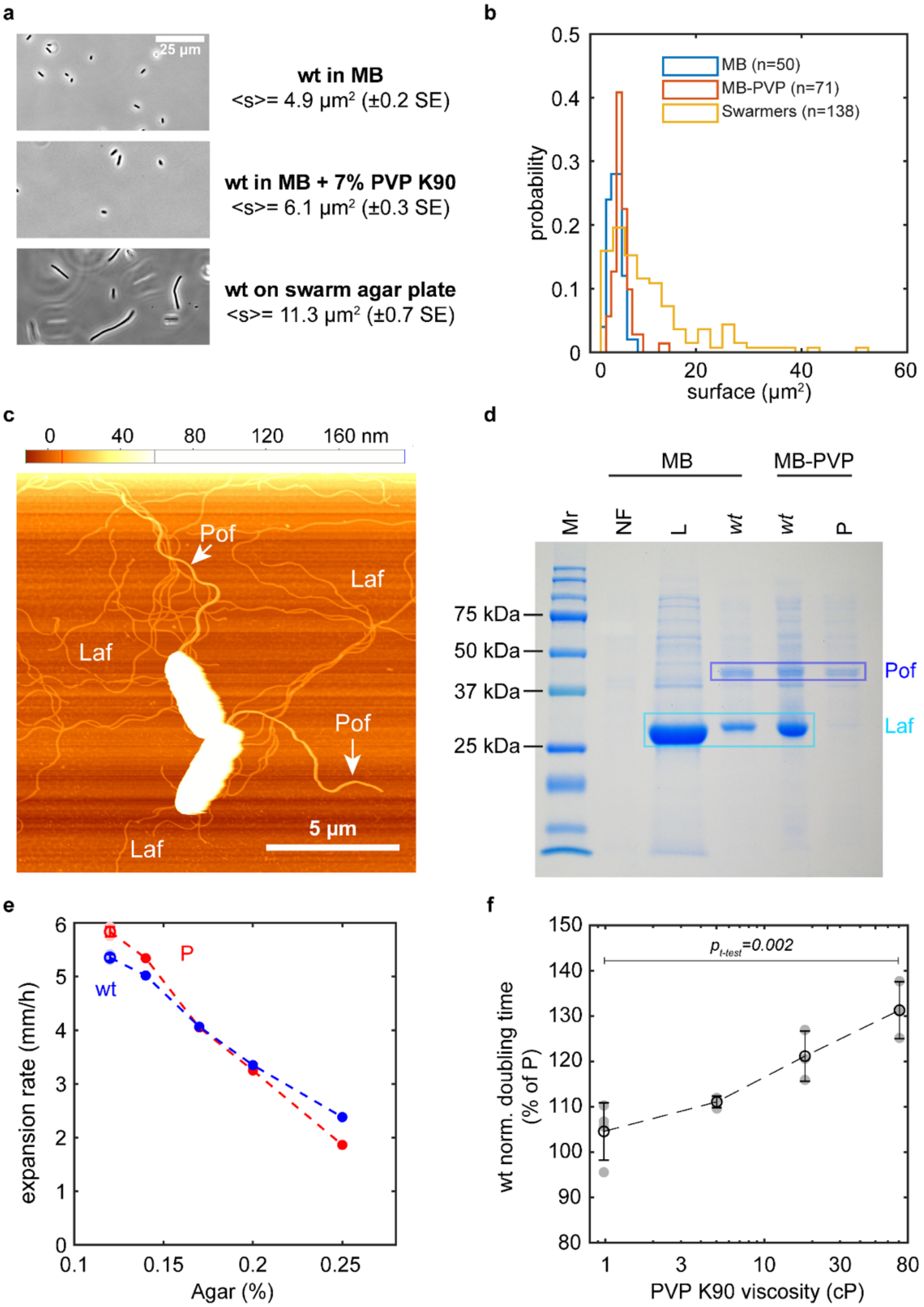

**S.I. Fig. 1 Phenotypic characterization of P and PL phenotypes and the cost of lateral flagella expression.** We established a characterization of the *V. alginolyticus* strains isolated by Kawagishi *et al*[2]. listed in Supplementary Table 3.

a-b) Size of wildtype cells grown under different conditions, as measured by the projected area of the cell body (b) from phase contrast microscopy images recorded at 40x magnification (a). While swarmer cells grown on semi-soft agar (1% agar/MB) approximately double in size compared to those grown in broth, those grown in viscous solutions (MB + 7% PVP K90) only show a minor increase in body length. The expression of lateral flagella which occurs both on surfaces and in viscous solutions is therefore not tightly coupled with body length.

c) AFM image of the wildtype strain grown in MB + 7% PVP K90, forming the PL phenotype, dried onto a glass slide. One polar flagellum (Pof) and multiple lateral flagella (Laf) can be observed. The polar flagella observed over several fields of view in this dried sample had an average height of 20.2 ± 1.4 nm (n=9), clearly distinguishable from the lateral flagella with 8.4 ± 1.5 nm (n=12) .

d) SDS-PAGE analysis of isolated flagella, for the wildtype, polar-only (P, YM4) and laf-only (L, YM19) strains grown in broth (MB) or viscous broth (MB + 7% PVP K90). The polar (Pof) and lateral (Laf) flagellins were identified by comparison between different strains and confirmed by size similarity with other species with dual flagellar architecture such as *S. piezotolerans*[3], *P. shigelloides*[4] and others[5]. The wild type strain expresses substantially more lateral flagella during growth in the presence of PVP, although some residual expression occurs in broth without PVP.

e) Ring expansion rate in soft agar MB plates at 30°C as a function of agar concentration for the wildtype (blue, 138-2) and polar-only (red, YM2) strains. Faint individual points at 0.12% agar are all from biologically independent experiments, with their average and SD displayed as a solid circle with error bars. Data at higher concentrations reflect a single experiment.

f) Doubling time of the wildtype strain as a function of viscosity of the growth medium, normalized by that of the Pof-only strain (YM4) at 26°C. The viscosity was altered by addition of PVP K90 to the MB growth medium, from 0% (0.98 cP at room temperature) to 7% (approximately. 70 cP). Each grey dot at a given viscosity represents an average across at least 5 technical replicates (wells) from one pair of bacterial cultures. Solid black circles and errors bars represent averages and SD across biological replicates. The slight growth deficit of the wildtype at 0% PVP may result from the residual expression of lateral flagella observed in (d).

**S.I. Figure 2**

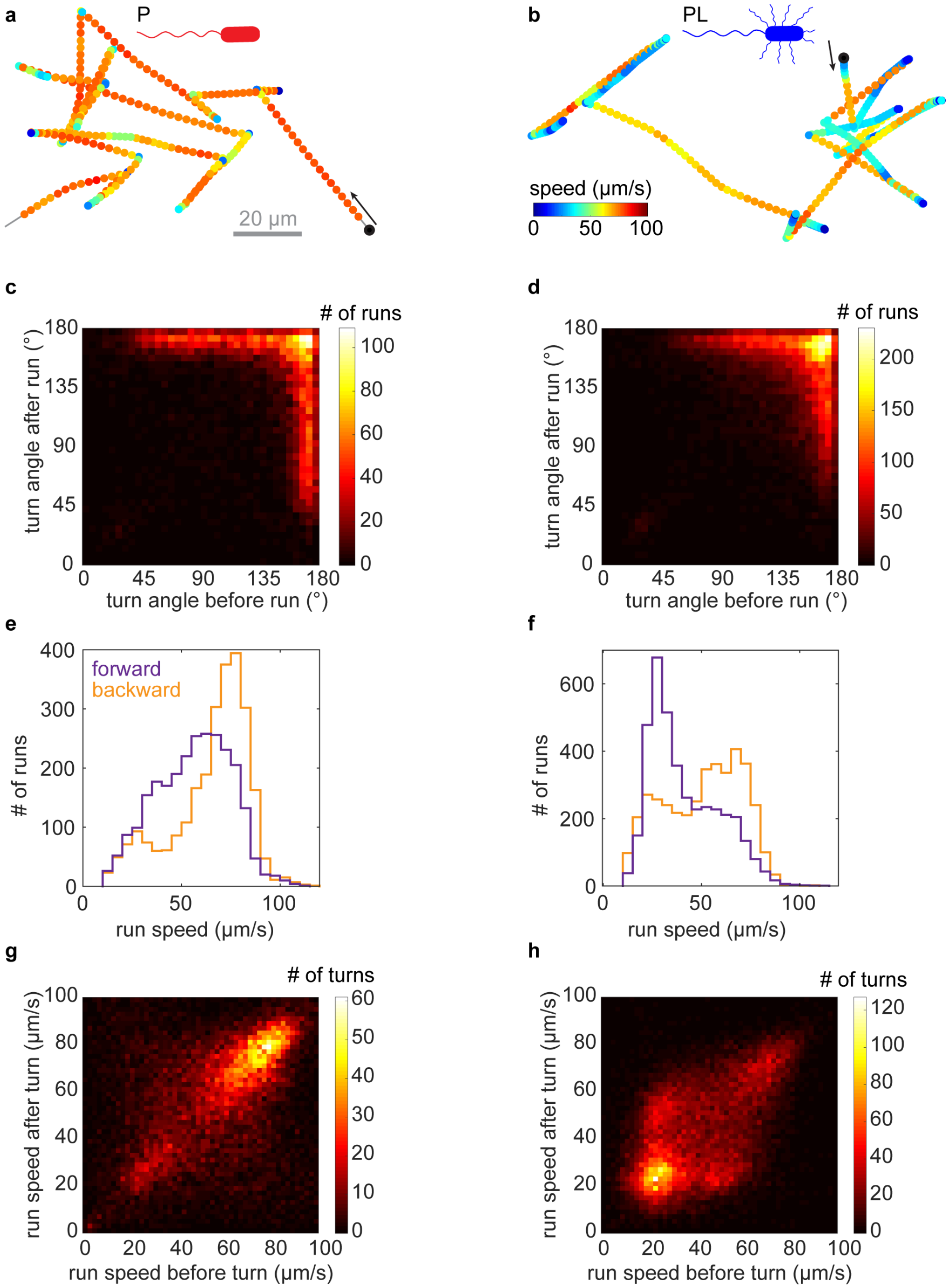

**S.I. Fig. 2: Run-reverse-flick motility in buffer.**

a, b) The same P (a) and PL (b) example trajectories as in Fig. 1a,b, but colored by instantaneous speed.

c,d) Bivariate distributions of turning angles preceding and succeeding a given run for the P (c) and PL (d) phenotype.

e,f) Histogram of average run speeds for forward (purple) and backward (runs).

g,h) Bivariate histograms of average run speeds before and after a turn for the P (g) and PL (h) phenotype. While successive runs have the same speed for the P phenotype (a), PL (b) additionally shows alternating fast and slow run speeds.
Data in panels a, b was recorded in motility chambers, while those in panel c-f was recorded in chemotaxis chambers with a 50 µM/mm L-serine gradient.

#### S.I. Figure 3

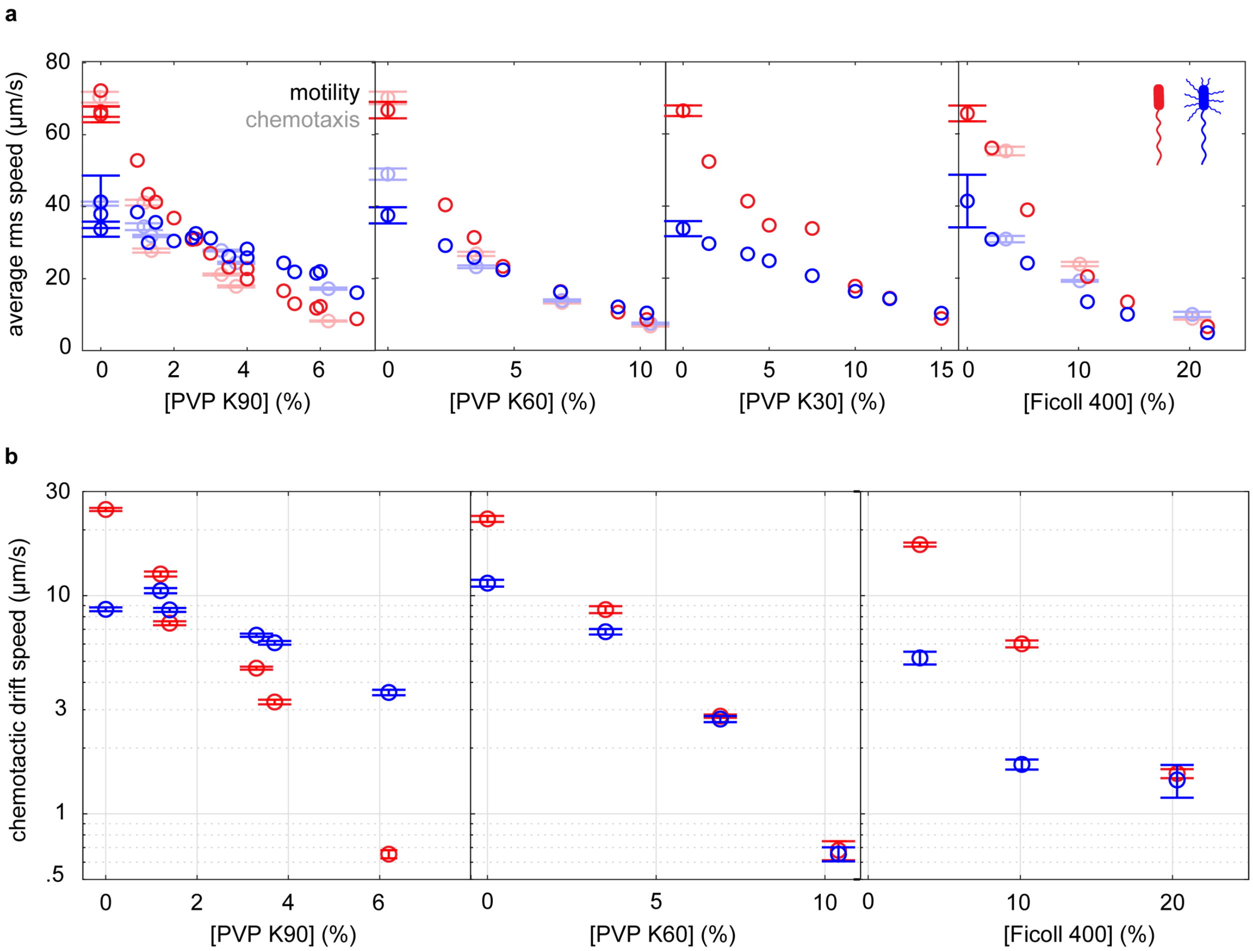

**S.I. Fig. 3: Swimming and chemotaxis in polymer solutions of varying concentrations.**

a) Average individual rms swimming speed measured for the P (blue) and PL (red) phenotypes in chemotaxis experiments (transparent shade, datasets 1-4) or motility chambers without gradients (solid shade, dataset 7), as a function of polymer concentrations in TMN. In motility chambers at 0% polymer, error bars represent standard deviations between technical triplicates, if available. In chemotaxis experiments, error bars represent the 95% confidence interval, computed from the standard errors of the mean which are estimated by jackknife resampling a single dataset into subsets consisting of 150 trajectories each.

b) Chemotactic drift speed of P (red) and PL (blue) phenotypes in a 50 µM/mm L-serine gradient as a function of polymer concentration in TMN. Error bars represent standard errors of the mean estimated by jackknife resampling a single dataset into subsets consisting of 150 trajectories each.

**S.I. Figure 4**

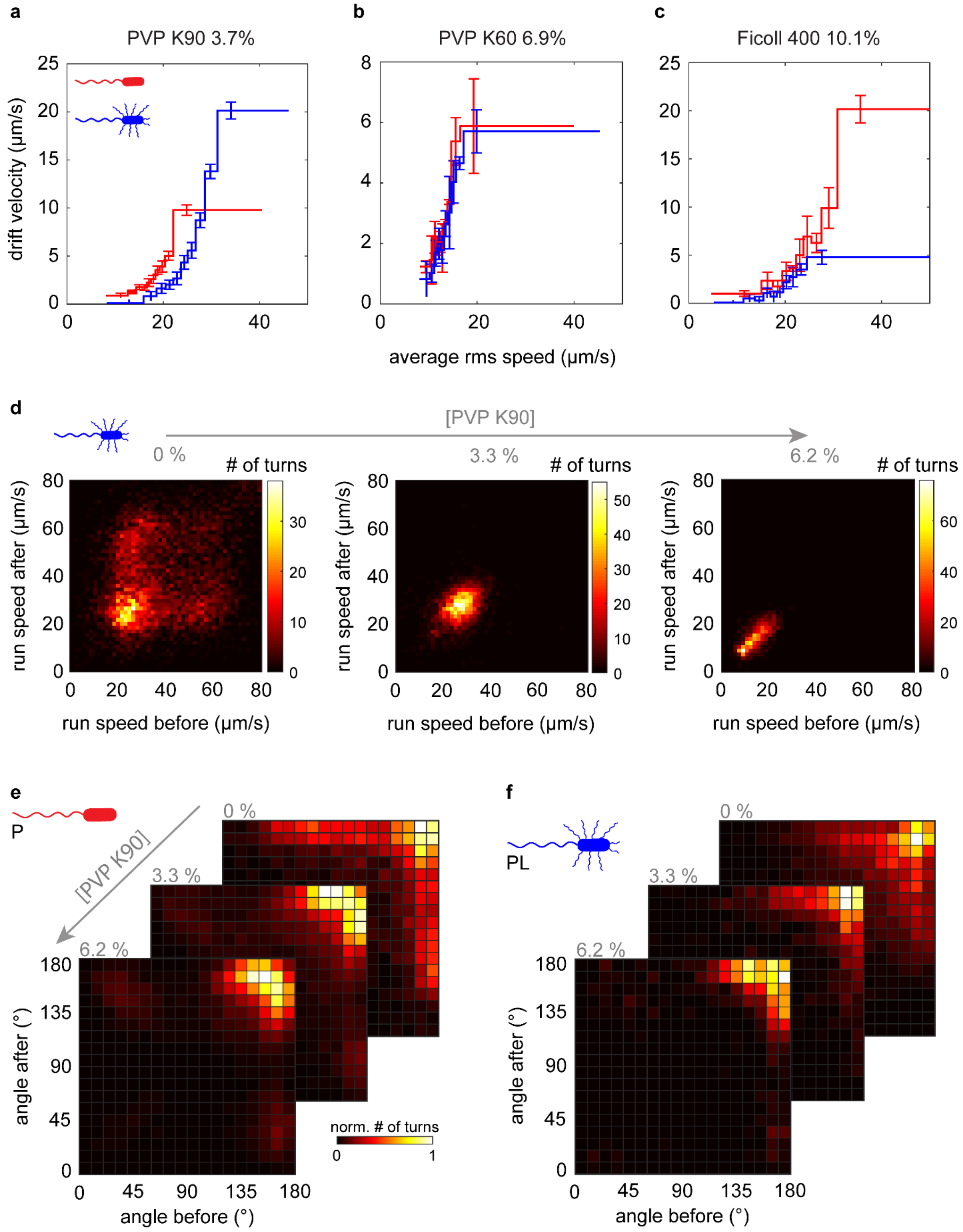
a
PVP K90 3.7%
b
PVP K60 6.9%
c
Ficoll 400 10.1%
drift velocity (µm/s)
average rms speed (µm/s)
d
[PVP K90]
0 %
3.3 %
6.2 %
### of turns
run speed after (µm/s)
run speed before (µm/s)
e
P
[PVP K90]
0 %
3.3 %
6.2 %
angle after (°)
angle before (°)
norm. # of turns
f
PL
0 %
3.3 %
6.2 %
angle after (°)
angle before (°)

**S.I. Fig. 4: Chemotactic performance and underlying behavior of PL and P phenotypes in polymer solutions.**

a-c) Chemotactic drift speed given average individual rms swimming speed in three different polymer solutions reveal no advantage of the PL over the P phenotype at fixed speed. For each flagellar architecture and medium, all trajectories of more than 1 s are distributed between 10 bins such that each bins contains trajectories of similar cumulative duration. The average drift velocity is computed for each bin. Error bars reflect 95% confidence intervals estimated by a jackknife resampling procedure.

d) Bivariate run speed distributions before/after a turn for the PL phenotype at increasing concentration of PVP K90, measured for the same culture. The alternating run speeds observed in buffer (0%, left) are quenched at 3.3% PVP K90 (middle) and 6.2% (right) in favor of an almost constant, slower run speed

e) Bivariate angular distributions before/after a run for the P and PL phenotypes at increasing concentration of PVP K90, showing that the motility pattern approximates run-reverse-motility at high concentrations for both. The small number of low-angle turns at increasing polymer density are likely false positives arising from increasing localization errors.

#### S.I. Figure 5

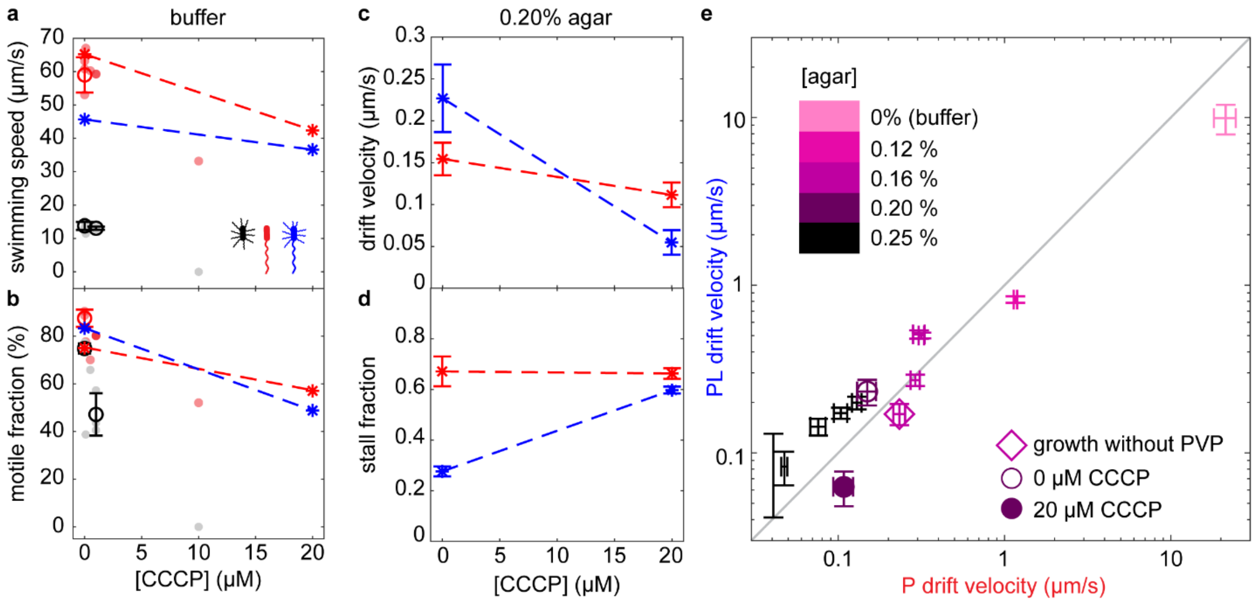

**S.I. Fig. 5. Active role of lateral flagella in chemotaxis in agar hydrogels.**

a,b) The average swimming speed (a) and motile fraction (b) of the PL (blue), P (red) or L (black) phenotypes swimming in buffer after a 60-min exposure to different concentrations of the protonophore CCCP (see Methods) which disrupts the proton gradient driving the rotation of lateral flagella. The L phenotype, corresponding to a Pof- mutant strain, is already completely non-motile at 10 µM CCCP, while the P and PL phenotypes maintain motility up to at least 20 µM CCCP. Faded dots are measurements conducted on the same cultures. Empty circles represent averages of technical replicates with error bars reflecting standard deviations between at least three. Asterisks connected by a dashed line are measurements performed on the same cultures as in panels c,d.

c,d) In a 50 µM/mm L-serine gradient in 0.20% agar, we measure the drift velocity (c) and stall fraction (d) of the P and PL phenotypes in conditions that do not (0 µM CCCP) or do (20 µM CCCP) disrupt the rotation of lateral flagella. Error bars reflect the standard error of the mean, estimated by a jackknife resampling procedure consisting of dividing the data into subsets of 300 trajectories and computing the SEM across subsets.

e) Drift velocity of the PL phenotype against the drift velocity of the P phenotype at a range of agar gel concentrations. Each point corresponds to one independent experiment where P and PL populations were studied in chemotaxis chambers prepared at the same time, except at 0% (buffer) where data from three biological replicates are combined. Error bars reflect SEM, estimated by a jackknife resampling procedure consisting of dividing the data into subsets of 300 (agar) or 150 (liquid buffer) trajectories and computing the standard error of the mean across subsets. As a control (diamond), both strains are grown without PVP to avoid the induction of lateral flagella in the wildtype strain. For 0.20% agar, experiments with the same cultures were performed in the absence (open circle) and presence (filled circle) of 20 µM CCCP.
